# Human skeletal muscle possesses both reversible proteomic signatures and a retained proteomic memory after repeated resistance training

**DOI:** 10.1101/2024.11.19.624068

**Authors:** Juha J. Hulmi, Eeli J. Halonen, Adam P. Sharples, Thomas M. O’Connell, Lauri Kuikka, Veli-Matti Lappi, Kari Salokas, Salla Keskitalo, Markku Varjosalo, Juha P. Ahtiainen

## Abstract

Investigating repeated resistance training separated by a training break enables exploration of the potential for a proteomic memory of resistance training (RT)-induced skeletal muscle growth. Our aim was to examine skeletal muscle proteome response to 10-week RT (RT1) followed by 10-week training cessation (i.e. detraining, DT), and finally, 10-week retraining (RT2). Thirty healthy, untrained participants conducted either periodic RT (RT1-DT-RT2, n=17) or a 10-week no-training control period (n=13) followed by 20 weeks of RT (n=11). RT included twice-weekly supervised whole-body RT sessions, and resting vastus lateralis biopsies were obtained every ten weeks for proteomics analysis using high-end DIA-PASEF’s mass spectrometry. The first RT period altered 150 proteins (93% increased) involved in e.g. energy metabolism and protein processing compared with minor changes during the no-training control period. The proteome adaptations were similar after the second RT compared to baseline demonstrating reproducibility in proteome adaptations to RT. Many of the proteins induced by RT1 were reversed towards baseline after detraining and increased again after retraining. These reversible proteins were especially involved in aerobic energy metabolism. Interestingly, several proteins increased after RT1 remain elevated after detraining, including carbonyl reductase 1 (CBR1) and proteins involved in muscle contraction, cytoskeleton and calcium-binding. Amongst the latter, calcium-activated protease calpain-2 (CAPN2) has been recently identified as an epigenetic muscle memory gene. We show that resistance training evokes retained protein levels even after 2.5 months of no training. This is the first study to demonstrate a potential proteomic memory of resistance training-induced muscle growth in human skeletal muscle.

**Key points:**

- Repeated resistance training in humans separated by a training break (i.e. detraining) enables the identification of temporal protein signatures over the training, detraining, and retraining periods as well as studying reproducibility of protein changes to resistance training.
- Muscle proteome adaptations were similar after a second period of resistance training when compared to baseline, demonstrating reproducibility in proteome adaptations to earlier resistance training.
- Many of the proteins induced by resistance training were reversed towards baseline after detraining and increased again after retraining. These reversible proteins were especially involved in aerobic energy metabolism.
- Several proteins increased after resistance training remain elevated after detraining, including carbonyl reductase 1 (CBR1) and calcium-binding proteins such as calpain-2 (CAPN2), a recently identified epigenetic muscle memory gene.
- Human skeletal muscle experiences retained protein changes following resistance training persisting over two months demonstrating a potential proteomic memory of resistance training-induced muscle growth.

Human skeletal muscle proteome response was investigated after 10-week resistance training (RT1) followed by 10-week training cessation (i.e. detraining, DT), and finally, 10-week retraining (RT2). Many of the proteins were reversed towards baseline after DT and increased again after RT2. These reversible proteins were especially involved in aerobic energy metabolism. However, several RT-induced proteins remain elevated after DT, including carbonyl reductase 1 (CBR1) and many proteins involved in muscle contraction or cytoskeleton and calcium-binding. Amongst the latter, calcium-activated protease calpain-2 (CAPN2) is a recently identified epigenetic muscle memory gene. This study shows that resistance training evokes retained protein levels even after 2.5 months of no training and demonstrates a potential proteomic memory of RT-induced muscle growth in human skeletal muscle. Created in BioRender.com.

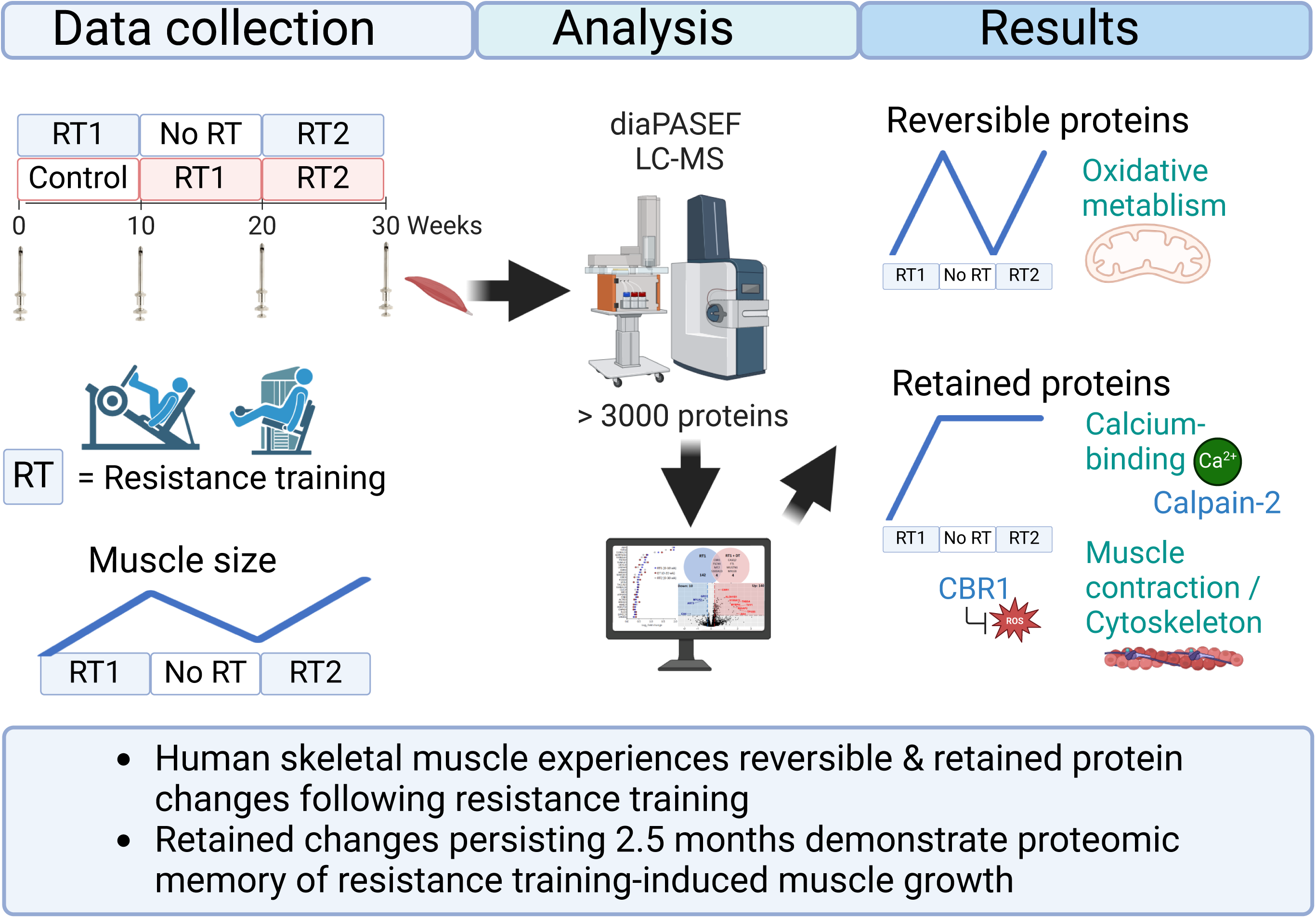

## Introduction

Resistance training (RT)-induced muscle hypertrophy and strength may be regained to an enhanced level after training breaks (i.e. detraining), a phenomenon termed muscle memory (Sharples & Turner, 2023). In a broader sense, muscle memory can also refer to a variety of long-term tissue-level effects of exercise. Thus, it can also potentially affect muscle maintenance and, therefore, an individual’s health and wellbeing, even in situations when there are breaks from exercise.

Previous (Egner *et al*., 2013) and more recent (Murach *et al*., 2020; Eftestøl *et al*., 2022) cellular and molecular muscle memory studies have concentrated on ‘cellular’ memory, i.e. retention of myonuclei during detraining in rodents. To date, human muscle memory and long-term adaptation studies to RT have primarily focused on either myonuclear retention (Psilander *et al*., 2019; Snijders *et al*., 2019; Cumming *et al*., 2024) or retained epigenomic and transcriptomic signatures in skeletal muscle of young adults (Seaborne *et al*., 2018) and older individuals (Blocquiaux *et al*., 2022). In addition, other types of exercise, such as high-intensity interval training (HIIT), have been investigated in humans (Pilotto *et al*., 2024) and progressive weighted wheel running in rodents (Wen et al., 2021) using a training, detraining and retraining design. Collectively, these studies suggest that there may be retention of myonuclei and epigenetic modifications, in particular, retained DNA methylation signatures from earlier training into detraining, leading to continued elevation of gene expression even during detraining. However, whether there are long-term or retained signatures following exercise training at the protein level is currently unknown.

Proteomics analysis, involving the quantification of thousands of proteins from a single muscle sample, can reveal complex adaptations to RT in skeletal muscle. Unfortunately, due to the high cost of proteomics analysis, many studies have had very small n-sizes, and/or the samples have been pooled decreasing statistical power. Nevertheless, a few recent proteomic studies have revealed that RT (Robinson *et al*., 2017; Du *et al*., 2024; Roberts *et al*., 2024; Moesgaard *et al*., 2024; Jessen *et al*., 2024) and other types of high-intensity exercise (Robinson *et al*., 2017; Deshmukh *et al*., 2021) can induce the remodelling of skeletal muscle at the individual protein level. Moreover, the effect of muscle disuse on muscle proteome has been investigated (Murgia *et al*., 2022). However, there is no evidence of long-term proteome adaptations following detraining after an earlier period of RT and how this affects protein changes to a later repeated period of RT. This poses the question of whether there is a memory of RT or RT-induced muscle growth evident at the protein level.

We hypothesized that RT would significantly and systematically affect the muscle proteome and that we would discover proteins with reversible (i.e. return to baseline levels during detraining) and some with retained signatures after detraining. However, due to the novelty of the study design and unbiased proteomic approach, we did not adopt an a priori hypothesis regarding which proteins or biological processes would be affected. Using this approach, we aimed to identify temporal protein signatures over the training, detraining, and retraining period, of which the retained proteins identified following earlier training may be implicated in skeletal muscle memory.

## Methods

### Study design and setting

This is a secondary analysis of a randomized and controlled study, of which the basic phenotype data and detailed description of study design, study participants, recruitment, and phenotyping methods have been recently published (Halonen *et al*., 2024). The study design is illustrated in Fig. 1. Briefly, participants were randomly assigned to periodic RT (PRT) or continuous RT (CRT) that included a control period of no-RT (Fig. 1). The PRT group completed a 10-week RT period (RT1), a 10-week detraining period (DT), and a repeated 10-week RT period (RT2) identical to RT1. The CRT group began with a 10-week non-RT period, followed by a 20-week continuous RT period.

**Figure 1.**
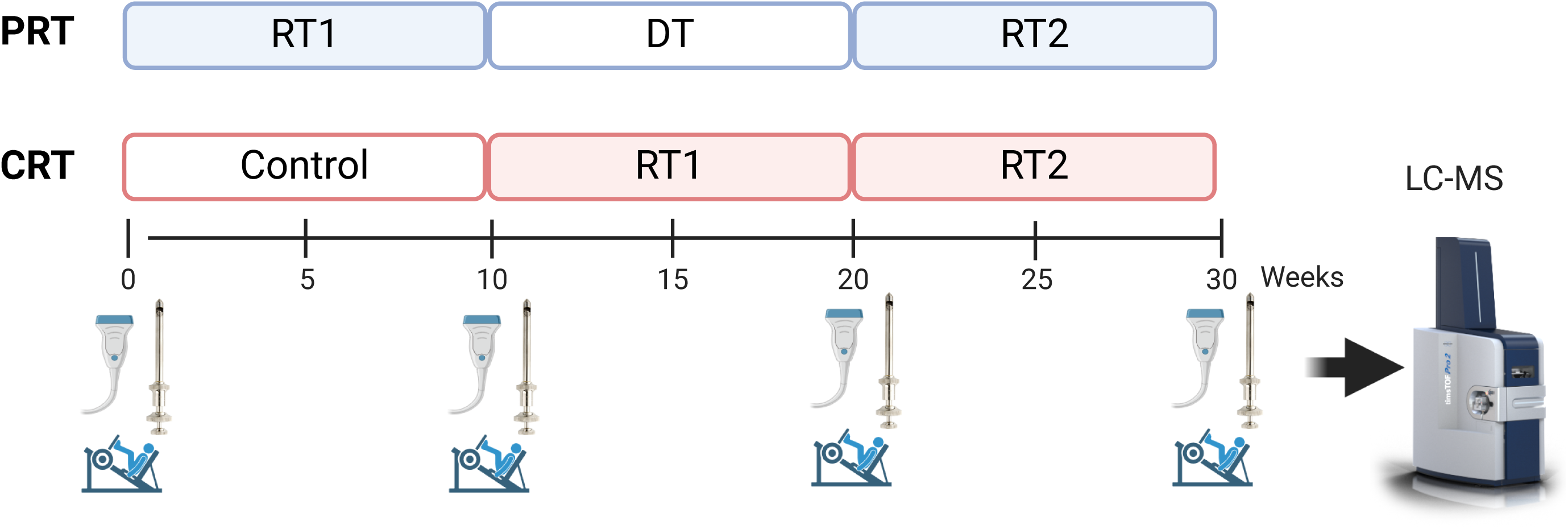
Study design. RT = resistance training. DT = detraining (no-RT). The periodic resistance training (PRT) group completed a 10-week initial RT period (RT1) followed by 10 weeks of no-RT/ detraining (DT) and then a second period of 10 weeks RT (RT2) with proteome analysed from all the participants n=17. The continuous resistance training (CRT) group completed a period of 10 weeks of no-RT (control) followed by continuous RT for 20 weeks (two periods of 10 weeks denoted RT1 & RT2 on figure) with n=13, of which n = 11 completed the training with available biopsies for proteomics.

### Participants

Fifty-five eligible males and females volunteered to participate in the original study of which forty-two participants completed the intervention (Halonen *et al*., 2024). Homogeneity between groups at baseline was achieved by separately dividing male and female participants into matched pairs by the combined Z score for BMI and age. For the present study, we included participants where we had at least 20 mg biopsy samples available for proteomics analysis resulting in a total of 30 participants. In the PRT group, which is the main focus of the present study, there were n = 17 (33.8 ± 5.2 years, 6 females/ 11 males), and in the CRT group n = 13 (31.8 ± 4.0 years, 6 females / 7 males) of which 2 (1 female/ 1 male) had samples only from the first time-points, i.e. before and after the control period). Inclusion criteria were: Age within the range 18-40 years, no regular RT history (< 10 RT sessions a year), not participating in systematic endurance-type training (< 2 endurance exercise sessions lasting > 30 minutes per week for the last six months), body mass index (BMI) within the range 18.5-30 kg/m^2^, and not currently consuming any anti-inflammatory drug(s). Exclusion criteria were: A history of medication that could affect exercise responses, use of creatine, any acute or chronic illness affecting cardiovascular, respiratory, musculoskeletal, and/or endocrine function, any other condition that may limit the ability to perform RT and testing (e.g., uncontrolled hypertension, diabetes, arthritic conditions, neuromuscular complications), or blood-borne diseases, diseases and medication affecting blood clotting, allergies to anaesthetic drugs, and severe psychological disorders. With these criteria and matching, there was no difference in the age (*P* = 0.295), height (*P* = 0.712), BMI (*P* = 0.657), leg press 1RM (*P* = 0.765), or VL CSA (*P* = 0.564) between the groups at the start of the intervention.

The study received ethical approval from the ethics committee at the University of Jyväskylä (857/13.00.04.00/2021). The study was conducted according to the declaration of Helsinki. All the applicants were informed of all potential risks of the study and had the possibility of withdrawing from the research project at any time, after which they signed an informed consent document. Personal data was stored and handled according to the ethical and GDPR guidelines of the University of Jyväskylä. The original study (Halonen *et al*., 2024) has been registered into ClinicalTrials.gov (ID: NCT05553769).

### Vastus lateralis muscle size and strength

The measurements and analysis have been explained in detail in (Halonen *et al*., 2024). Briefly, Vastus lateralis (VL) muscle cross-sectional area (CSA, cm^2^) was measured using a B-mode axial plane ultrasound (model SSD-α10, Aloka, Tokyo, Japan), which has a high correlation with MRI in detecting changes in muscle CSA (Ahtiainen *et al*., 2010). The ultrasound images were analysed using ImageJ software (version 1.54). The coefficient of variation percentage (CV%) for VL CSA from test to retest was 1.6%. One participant from the PRT group was excluded from the VL CSA analysis due to poor image quality, however this participant was included in all the other analyses.

### RT intervention

Both groups underwent the two identical 10-week RT periods, but the difference between the groups was that the CRT group performed the first and second 10-week RT periods continuously, and the PRT group had a 10-week detraining period between the two RT periods (Fig. 1). The twice-weekly exercises were leg press (4 x 8-10 reps), knee extension (4 x 8-10 reps), Smith machine bench press (3 x 8-10 reps), barbell biceps curl (4 x 8-10 reps), and chest supported seated row (4 x 8-10 reps) with two-minute rest periods between the sets. The participants were instructed to perform 8-10 reps in each set with approximately 1-2 repetitions in reserve, except in the final set of each exercise in the second training session of every week, when the last set was performed until volitional failure. To enable progression, the number of repetitions was then used to adjust the training loads for the following week; if the number of performed repetitions was more than 10, the load was increased, and if the repetitions were less than eight, the load was decreased. A more detailed description of the training is explained in (Halonen *et al*., 2024). All training sessions were performed at the laboratory of the University of Jyväskylä’s Faculty of Sport and Health Sciences, and the sessions were instructed and supervised to ensure correct exercise techniques.

### Control period and detraining

During the control period in the CRT group and after the first 10-week RT period in the PRT group (start of DT), the participants were instructed to continue their habitual lifestyle but to avoid any form of resistance or endurance-type training or any other unaccustomed exercise for the next 10- week period. Physical activity and other lifestyle changes during the detraining period were assessed with a survey (Halonen *et al*., 2024) and the participants were also contacted via email in the middle of the detraining period to ensure that they did not participate in any form of RT. These analyses suggested there was no RT conducted during the detraining period (Halonen *et al*., 2024).

### Muscle biopsy obtainment

The muscle biopsies were obtained at 0, 10, 20, and 30 weeks, at rest ∼6 days after the last exercise bout, except in the CRT group after 10 weeks of RT1, where the biopsy was taken ∼3 days after the last exercise bout to minimize training break in the middle of the continuous 20-week RT intervention. Biopsies were taken from vastus lateralis 1/3 of the length from the patella towards the greater trochanter with local anaesthetic and a 5-mm Bergström biopsy needle together with suction as previously described (Hulmi *et al*., 2009). The biopsy was always from the right leg. The second biopsy was taken approximately 1 cm medially to the first biopsy, the third biopsy 1 cm laterally to the first biopsy, and the fourth biopsy 1 cm medially to the second biopsy. The muscle sample was cleaned of any visible connective tissue, adipose tissue, and blood and frozen immediately in liquid nitrogen (-180 °C) and stored at -80 °C.

### Sample preparation for proteomics analysis

Twenty mg of muscle tissue was disrupted and lysed using “high” setting with Beatbox (PreOmics, Germany) using Tissue kit 24x (P.O.00128, PreOmics). The total protein concentration of the homogenates was measured with Bio-Rad Protein Assay Dye (#5000006, Bio-Rad Laboratories). The samples were stored frozen at -80°C overnight followed by digestion. 50 µg of total protein from each sample was taken for proteomics analysis. Sample digestion and purification were done on all samples at the same time with an iST HT kit (P.O.00067, PreOmics) according to the manufacturer’s instructions. After desalting and purification, the samples were dried in a centrifuge concentrator (Concentrator Plus, Eppendorf). The dried peptides were reconstituted in 30 µl buffer A (0.1% (vol/vol) TFA, 1% (vol/vol) acetonitrile (#83640.320, VWR) in HPLC grade water (#10505904, Fisher Scientific)). For the DIA analysis, the resuspended peptides were further diluted 1:50 in buffer A1 (1% formic acid in HPLC water). Twenty µl was loaded into an Evotip (Evosep, Denmark) according to the manufacturer’s instructions and run at the same time.

### Mass spectrometry and data analysis

The samples were analysed using the Evosep One liquid chromatography (LC) system coupled to a hybrid trapped ion mobility quadrupole TOF mass spectrometer (Bruker timsTOF Pro2, Bruker Daltonics) (Meier *et al*., 2018) via a CaptiveSpray nano-electrospray ion source (Bruker Daltonics). An 8 cm × 150 µm column with 1.5 µm C18 beads (EV1109, Evosep) was used for peptide separation with the 60 samples per day method (21 min gradient time). Mobile phases A and B were 0.1 % formic acid in water and 0.1 % formic acid in acetonitrile, respectively. The MS analysis was conducted in the positive-ion mode with dia-PASEF (data independent analysis, dia; Parallel Accumulation Serial Fragmentation, PASEF) method (Meier *et al*., 2020) with sample optimized data dia scan parameters. We performed data-dependent acquisition (DDA) in PASEF mode from a pooled sample to be able to adjust dia-PASEF parameters optimally. To perform sample-specific dia-PASEF parameter adjustment, the default dia-short-gradient acquisition methods were adjusted based on the sample-specific DDA-PASEF run with the software “tims Control” (Bruker Daltonics). The following parameters were modified for each sample type: m/z range: 415.0 – 1215.0 Da; mass steps per cycle: 32; mean cycle time: 1.27 s. The ion mobility windows were set to best match the ion cloud density from the sample type-specific DDA-runs.

To analyse diaPASEF data, the raw data (.d) were processed with DIA-NN v1.8.1 (Demichev *et al*., 2020, 2022) utilizing spectral library generated from the UniProt human proteome (USP000000589, downloaded 21.08.2023 as a FASTA file, 20159 proteins). During library generation, the following settings were used with fixed modifications: carbamidomethyl (C); variable modifications: acetyl (protein N-term), oxidation (M); enzyme:Trypsin/P; maximun missed cleavages: 1; mass accuracy fixed to 1.5e-05 (MS2) and 1.5e-05 (MS1); Fragment m/z set to 200-1800; peptide length set to 7- 50; precursor m/z set to 300-1800; Precursor changes set to 2-4; protein inference not performed. All other settings were left to default.

### Statistical and bioinformatics analysis of the proteomics data

The input file to further DIA data analysis was the DIA-NN Report.pg_matrix. For data pre- processing, an in-house R-script was utilized. Raw intensity values were log2 transformed and median-normalized. Afterward, missing values were imputed using QRILC imputation (https://CRAN.R-project.org/package=imputeLCMD). For time comparison, p-values were calculated with paired sample t-test and groups with Student t-test using python package SciPy (Virtanen *et al*., 2020), and adjusted using Benjamini-Hochberg (false discovery rate, FDR) method via statsmodels package (Seabold & Perktold, 2010). For the analysis, stringent *FDR* < 0.05, Fold change (FC) > |1.2| filtering was used if not otherwise mentioned. ShinyGO (http://bioinformatics.sdstate.edu/go/) (Ge *et al*., 2020) v0.81 overrepresentation analysis was used to identify enriched Gene Ontology (GO) terms from the GO (biological processes (BP), cellular components (CC), molecular function (MF) and Reactome (Jassal *et al*., 2019) databases. We used an FDR cutoff of 0.05 and the quantified proteome as background as recommended (Timmons *et al*., 2015; Wijesooriya *et al*., 2022). Categorical annotations for each protein in the form of GO terms of BP, CC, and MF were extracted from the UniProt database. The pathway entitled Metabolism was omitted as a too generic term from these figures. To show the connections between the significant pathways and the individual proteins, Sankey diagram was also created in R as were the Volcano plots.

All other figures were made with GraphPad Prism software (version 10.0, GraphPad Software Inc), Microsoft Excel (version 2410), BioRender.com, or using Proteomaps approach (Liebermeister *et al*., 2014). Part of the Venn diagrams were generated using: https://bioinformatics.psb.ugent.be/webtools/Venn/. In line or bar graphs, data is presented as mean ± standard error of the mean (SEM) by applying the inverse log_2_ transformation to estimate the original values of the proteins. The statistics were, however, always conducted from log_2_ values. In Proteomaps each protein is represented by a polygon of which size corresponds to the magnitude and consistency of the change after RT (i.e. fold change * -Log10(FDR)). It also situates functionally related pathways in adjacent regions based on KEGG Orthology identifiers. To plot temporal changes in proteins across the intervention, we undertook Self Organising Map (SOM) profiling of the change in standardised mean protein content within each time-point using Partek Genomics Suite as previously described earlier (Turner *et al*., 2020). SOM analysis plots a standardized mean (shifts mean to a value of 0 and scales to a standard deviation of 1) for the group of samples across each time-point to identify temporal signature changes over the time-course of the intervention.

## Results

### Changes in muscle size and strength

In this subcohort of our study (Halonen *et al*., 2024), the first 10 weeks of RT increased muscle CSA in the VL and 1 RM strength in the leg press (*P* < 0.001, Table 1). Moreover, when comparing periodic RT (PRT) and continuous RT (CRT) during the whole intervention (weeks 0 to 30), no significant differences were observed in muscle size or strength (Table 1). This occurred because the muscle size and strength lost during detraining (*P* < 0.001) were quickly regained during retraining in the PRT group. In contrast, the CRT group experienced a somewhat slower rate of gains in muscle size and strength during the second 10-week RT period (Halonen *et al*., 2024).

**Table 1.** Characteristics of the participants analysed for proteomics (Mean ± SD).

| Group | $\Delta\%$ VL CSA | | | $\Delta\%$ 1 RM | | |
| --- | --- | --- | --- | --- | --- | --- |
|  | (0-10 wk) | (0-20 wk) | (0-30 wk) | (0-10 wk) | (0-20 wk) | (0-30 wk) |
| PRT | 16.0 ± 8.5*** | 3.9 ± 7.7*** | 19.4 ± 9.5 | 20.8 ± 13.3*** | 15.4 ± 10.6 | 29.7 ± 16.2 |
| CRT | 1.3 ± 4.4 | 23.2 ± 10.1 | 27.4 ± 11.9 | 1.6 ± 4.4 | 22.2 ± 11.1 | 28.6 ± 12.3 |

1 RM strength measured in leg press. N = 16 in PRT except in the 1 RM (n=17) and n=11 analysed for proteomics in the CRT-group (except n=13 at 0-10 wk). \*\*\* $P < 0.001$ difference between PRT and CRT. The data with larger n-size including also other time-point comparisons has been published earlier (Halonen *et al.*, 2024).

### RT affects on the muscle proteome

Using an FDR cutoff of 0.01 on both peptide and protein levels, we identified 3610 proteins of which 3391 were unique. This led to the quantification of 3127 and 3130 unique proteins in PRT and CRT groups, respectively. The first RT period (RT1) in PRT significantly (*FDR* < 0.05, FC >|1.2|) altered 150 proteins compared to only three proteins in the no-RT control period (Fig. 2A). See all the time-point comparisons in Supporting information, Supplement 1 including all FDR-values and fold changes. The large majority 93% (140) of the total 150 proteins identified had increased levels after RT1. Within cellular processes and pathways, the altered proteins belonged to e.g., aerobic and glycolytic metabolism, amino acid metabolism, cytoskeleton, chaperones and protein processing (Fig. 2B).

**Figure 2.**
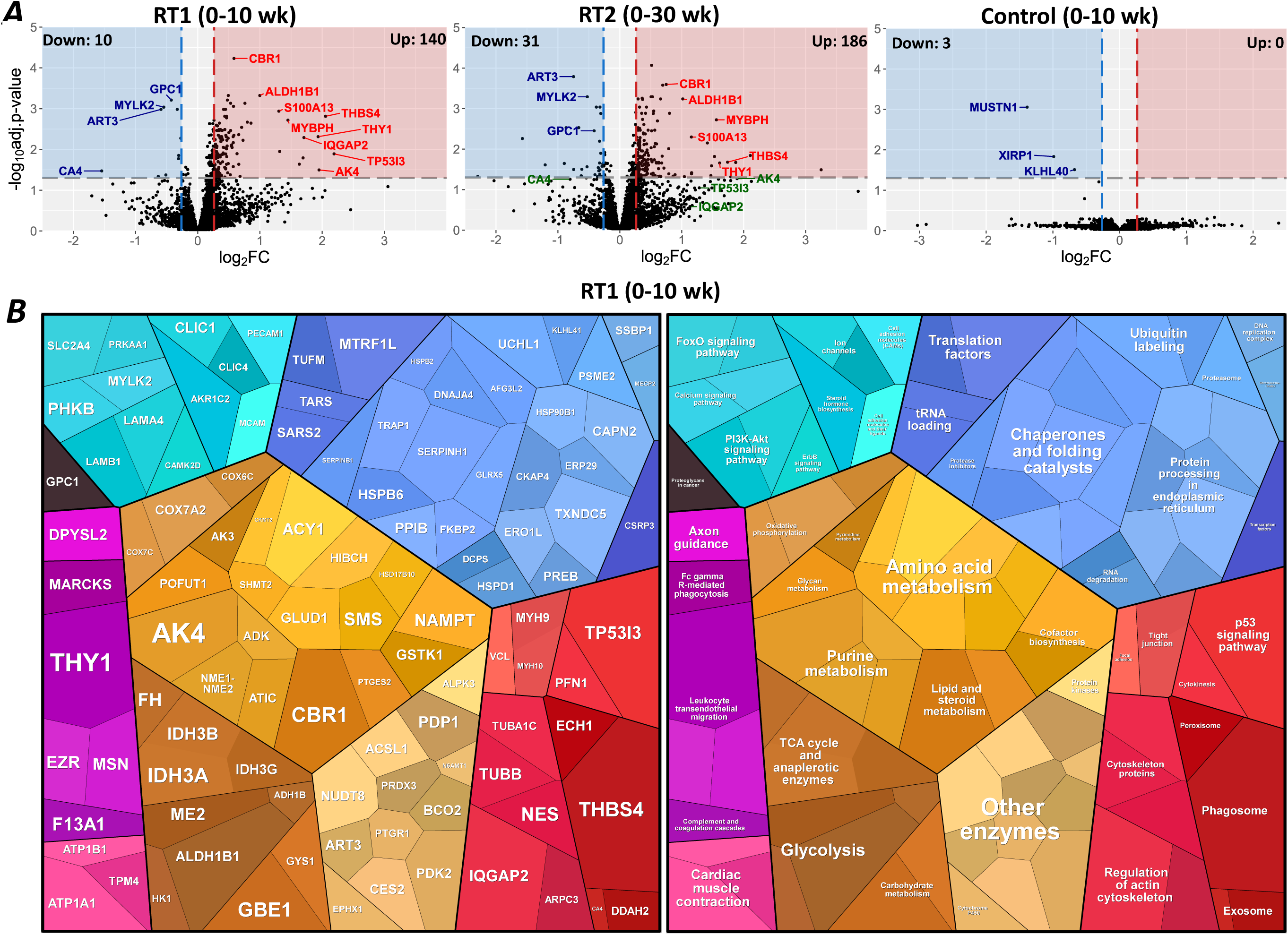
*A*, Volcano plot of the significantly altered proteins after the first 10-week RT (RT1) and RT2 in the PRT condition (n =17) and control (no RT) in the CRT condition (n=13). The grey line represents the significance cutoff (adj.p-value, i.e. *FDR* < 0.05: -log_10_ of 0.05 is ∼1.30) and FC >|1.2|: log_2_ of 1.2 is ∼0.26) and the significantly altered proteins are in the coloured boxes. Highly altered (highest log2FC * -log10(FDR)) proteins are labelled. In the RT2 plot, the same proteins are labelled as in the RT1 plot, while the green colour denotes those proteins that were significantly altered in RT1 but not in RT2. *B*, Proteomaps of the RT1 period in the PRT group. The proteins in the left panel are mapped in the right panel to functional KEGG categories. n=17 in PRT, n=11 in CRT and n=13 in the control period (CONT).

Next, we examined the changes following RT2 and whether these were similar to changes after the earlier RT1 period. When RT2 was compared to baseline, 217 proteins were significantly altered (186 / 86% increased, Fig. 3A, Supporting information, Supplement 1). Most of the proteins (66.6% or 100/150) altered after RT1 were also significantly altered after RT2 and abundance changes correlated between RT1 and RT2 (Fig. 3B). Also, the top upregulated proteins (Fig. 2A and 3C) were similar after RT1 and after RT2 (vs. baseline). This was confirmed further, as no proteins were significantly altered when post RT2 and RT1 timepoints were compared. However, there were 117 unique proteins significantly altered after RT2 (but not after RT1), with these proteins enriched in aerobic metabolism pathways and processes (Fig. 3D). When post-RT2 was compared to pre-RT2 (post-DT), only 17 proteins were significantly altered (Fig. 3A). However, half of these were shared in both RT1 and RT2, and the direction and overall magnitude of the responses were very similar (Fig. 3E).

**Figure 3.**
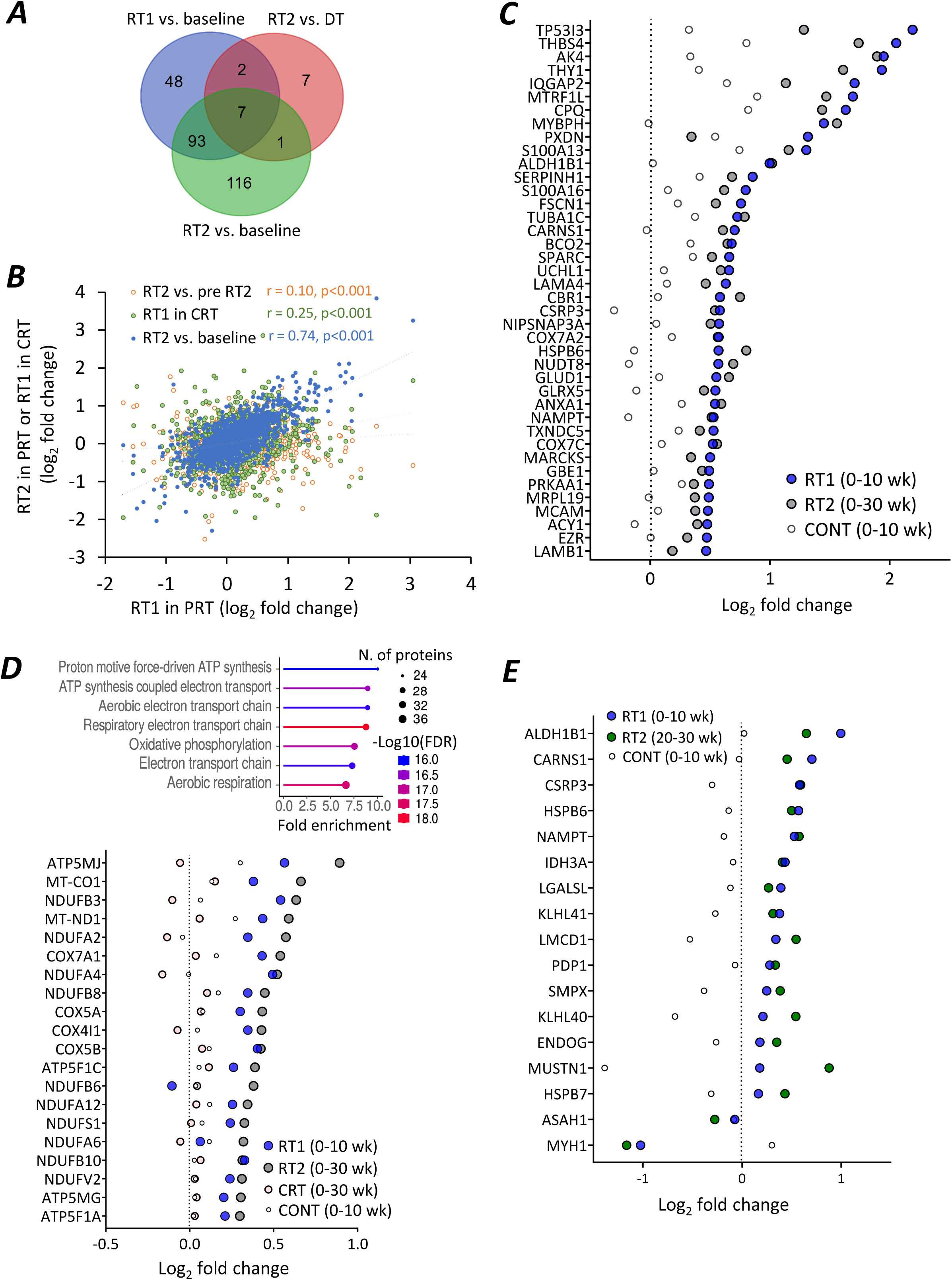
*A*, Venn Diagram analysis of the statistically differentially (*FDR* < 0.05, FC >|1.2|) regulated proteins following RT1 and RT2 compared to baseline (RT1+RT2) and RT2 vs. pre RT2 (RT2), illustrating the number of common overlapping and unique significant proteins across comparisons in the PRT group. *B*, Pearson’s correlation of protein abundance changes in different comparisons (in PRT between RT1 and RT2 vs. baseline, between RT1 in PRT and RT1 in CRT, and in PRT between RT1 and RT2 vs. DT (pre-RT2)). *C*, Top 40 upregulated proteins (*FDR* < 0.05) in PRT group after RT1 (vs. baseline) and changes in the same proteins during RT2 (vs. baseline). *D*, GO biological processes in the 117 proteins unique to RT2 vs. baseline illustrated by ShinyGo. The graph below figure shows that the top 20 of those RT2-induced proteins have not yet significantly increased after RT1. *E*, Altered proteins (*FDR* < 0.05 and FC> |1.2|) in RT2 vs. DT (pre-RT2) comparison and respective protein changes in RT1 in the PRT group and control period in the CRT group. n=17 in PRT, n=11 in CRT and n=13 in the control period (CONT). The exact *FDR*-values and fold changes of the proteins in each comparison are in Supporting information, Supplement 1.

### Temporal protein signatures identifies both reversible and retained proteins after RT

Proteome signatures in the PRT group were further analysed by self-organizing map (SOM) profiling. For the SOM analysis, we identified 285 proteins that were significant in at least one of the possible six comparisons between the time-points (Fig. 4A). It highlighted four predominant temporal signatures (Fig. 4B). Signature 1 included the smallest proportion of proteins (14.0%) that were decreased after RT. Signature 2 included proteins (20.3%) that were increased after RT1, fully returned to baseline, and then increased again after RT2. Signature 3 included proteins (37.2%) that were increased after RT1 and then returned towards, but not fully to baseline levels after DT, and then once again increased after RT2. Finally, signature 4 included proteins (28.4%) increasing after RT1 and then retained as elevated after DT and RT2. In addition, so that researchers can check the temporal profile of their protein of interest across RT1, DT, and RT2, we conducted SOM analysis also using all detected proteins (with no statistical or fold change cut-offs) and included them as a resource in Supporting information, Supplement 2.

**Figure 4.**
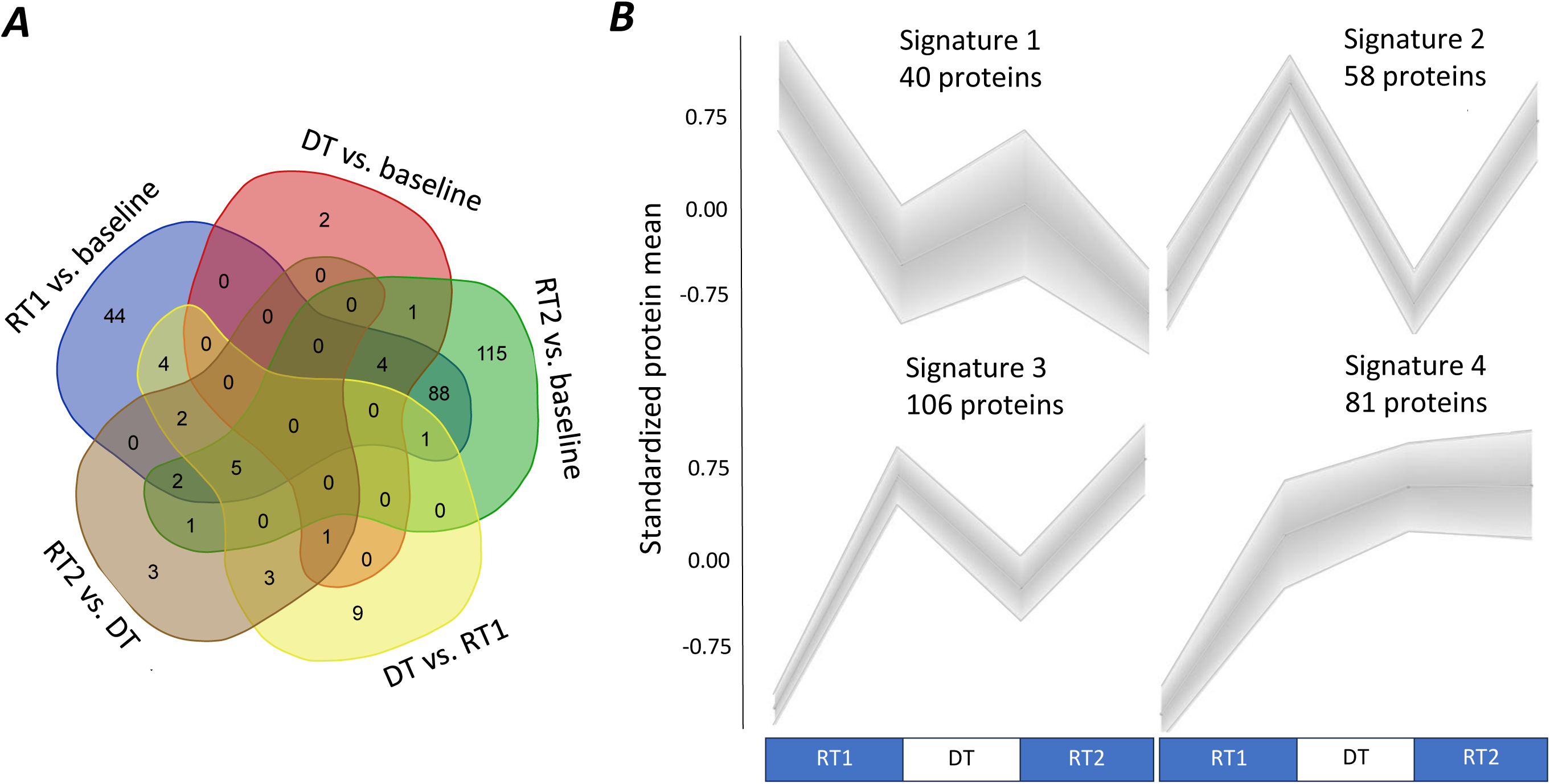
*A*, Venn diagram analysis of the statistically differentially regulated proteins: 1) RT1 vs. baseline, 2) DT vs. baseline, 3) RT2 vs. baseline, 4) DT vs. RT1 (pre-DT) and 5) RT2 vs. DT (pre-RT2) (*FDR* < 0.05, FC >|1.2|). The final 6th comparison: RT2 vs. RT1 is not shown as it did not produce any statistically significantly different proteins. The figure illustrates the number of common overlapping and unique significant proteins across each comparison. There were 285 proteins that were significantly different in at least one of the pairwise comparisons highlighted in the Venn diagram. *B*, SOM profiling of 285 proteins identified in the Venn diagram above in the PRT group over the 30 weeks time-course of baseline, RT1, DT and RT2. See also Supporting information, Supplement 2

### Proteins with reversible signature after RT1

Many proteins changed in response to RT1 and then returned towards baseline during detraining (signature 2 defined above). To investigate this protein signature further and to confirm that protein levels were back to baseline after detraining, we first identified the proteins that significantly increased after RT1. From those proteins, we selected the ones that demonstrated greater than a mean decrease of 70% after DT (DT vs. RT1 decrease > 70 %), thus confirming a large magnitude decrease after the DT-period. We further confirmed that these protein levels after DT were not different from baseline (FDR > 0.5 at post DT vs. baseline indicating high probably of no difference). With these criteria, we found 42 proteins, of which 36 were confirmed to belong to signature 2 (containing 58 proteins) and illustrated in Fig. 5A. These 36 reversible proteins are involved in aerobic energy metabolism (Fig. 5B). Two examples of completely reversible RT-upregulated proteins were kelch-like family protein KLHL41 recently identified as RT-induced protein (Jessen *et al*., 2024) and nicotinamide phosphoribosyltransferase (NAMPT), which is a rate-limiting enzyme in energy metabolism and nicotinamide adenine dinucleotide (NAD^+^) production, a coenzyme essential for muscle (Beltrà *et al*., 2023) (Fig. 5C). Indeed, in molecular function within GO terms reversible proteins were especially enriched in oxidoreductase and dehydrogenase functions and NAD (Fig. 5D).

**Figure 5.**
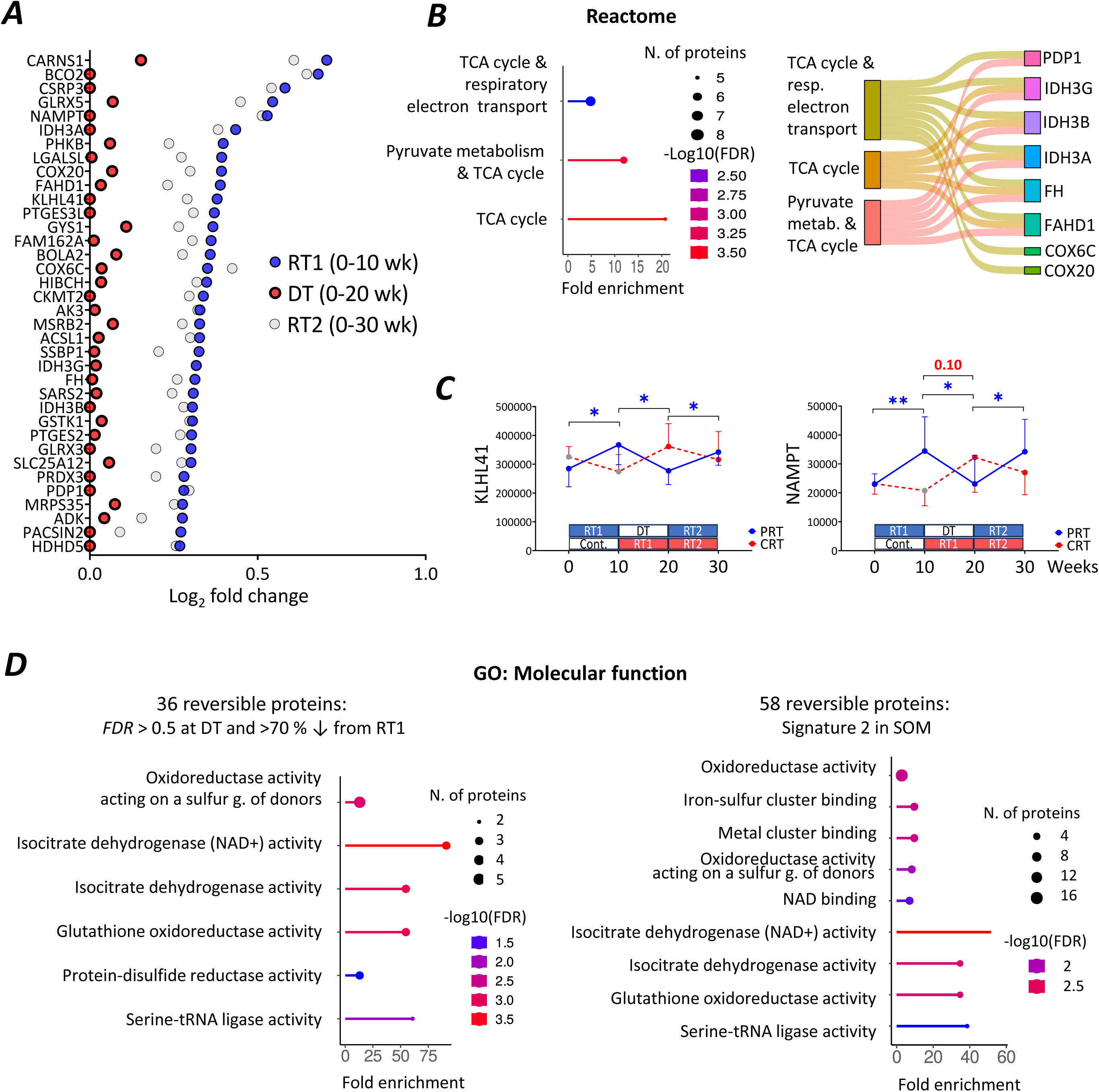
*A*, 36 proteins reversible proteins, which increased after RT1 in the PRT group and restored to the baseline after DT with the following criteria of: 1) *FDR* > 0.5 at post RT+DT vs. baseline, 2) relatively decreased > 70 % from the post-RT1, i.e. DT vs. RT1, and 3) belong to signature 2 (see Figure 4). *B*, Reactome pathway analysis together with Sankey-diagram show that reversible proteins belong to aerobic energy metabolism processes. *C*, Two examples of reversible proteins, KLHL41 and NAMPT. The proteins are plotted by applying the inverse log_2_ transformation to estimate their original mean ± SEM values. The statistics were conducted from log_2_ values. * <0.05, ** <0.01 or exact FDR-value (if *FDR* < 0.10) vs. baseline. *D*, Molecular function roles of reversible proteins. g. = group. N=17 in PRT and n=11 in CRT except n=13 in the first 10 weeks (control period). Figures *B* and *D* were extracted and modified from ShinyGO. The exact *FDR*-values and fold changes of the proteins in each comparison are in Supporting information, Supplement 1.

### Proteins with retained or ‘memory’ signatures after RT1, DT, and RT2

After DT, we found eight proteins altered compared to baseline and of those proteins, four were also increased after RT1 (*FDR* < 0.05, FC > 1.2, Fig. 6A). Three of these four proteins were contained within SOM profiling signature 4, i.e. increased after training and retained after detraining, including carbonyl reductase (CBR1), NAD-dependent malic enzyme (ME2) and protein S100-A13 (S100A13), whereas fascin (FSCN1) was contained within SOM signature 3 (Fig. 6B). Looking at the time-course within CRT and PRT groups reveals that out of these proteins (CBR1, ME2, S100A13 and FSCN1), especially CBR1 is an RT-induced "memory" protein as it is consistently increased by RT and staying elevated following DT and RT2 with very stable levels during the control period (Fig. 6B).

**Figure 6.**
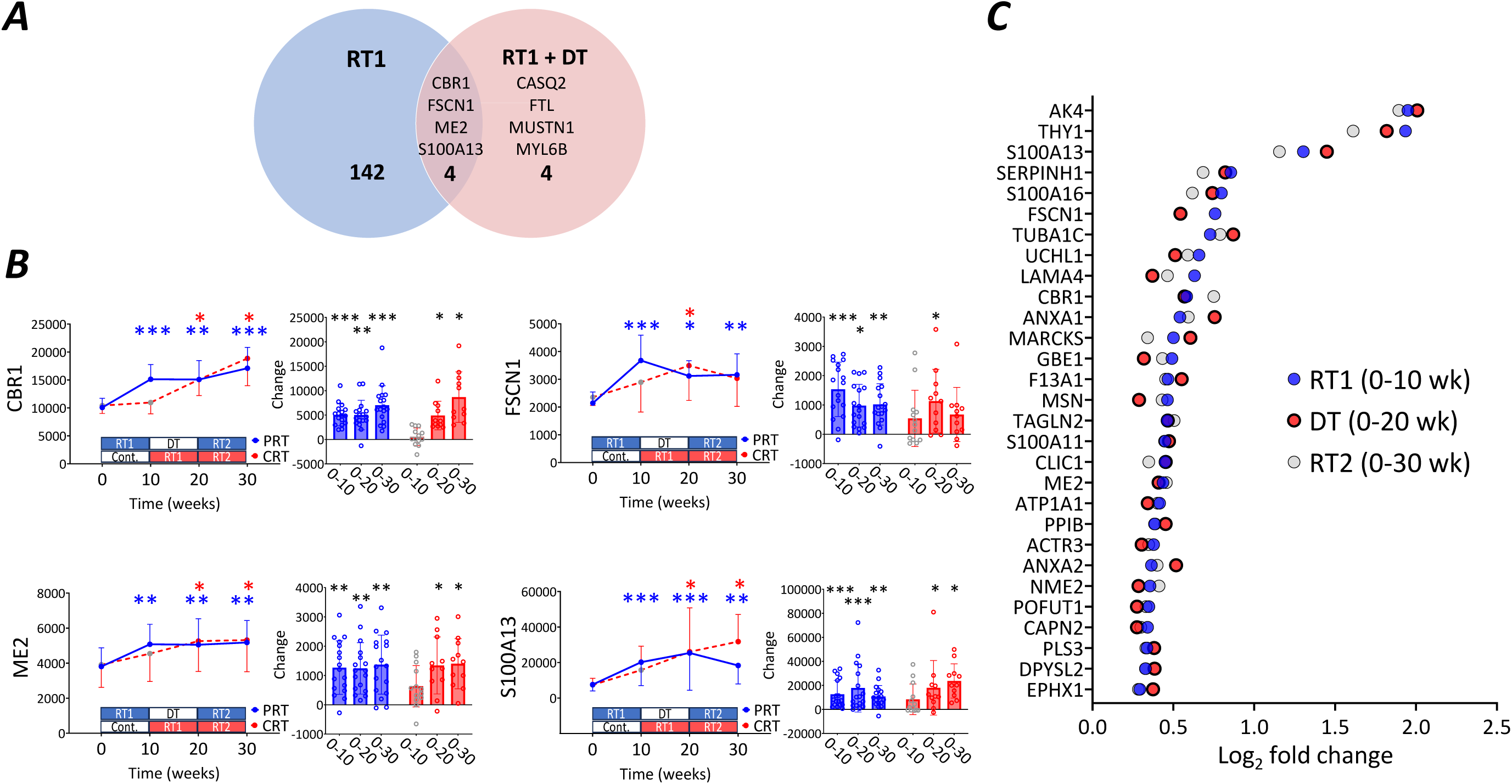
*A*, Venn Diagram analysis of the statistically differentially regulated proteins following RT1 and RT1+DT compared to baseline illustrating the number of common overlapping and unique significant proteins across comparisons. *B*, Four proteins increased after RT1 and retained as increased after DT (and RT2) at *FDR* < 0.05, FC >|1.2|. *C*, 29 proteins, which significantly increased after RT1 in the PRT group and remained elevated after DT and post RT2 (*FDR* < 0.1 and FC > 1.2). N=17 in PRT and n=11 in CRT except n=13 in the first 10 weeks (control period). The proteins are plotted by applying the inverse log_2_ transformation to estimate their original mean ± SEM values. The statistics were conducted from log_2_ values. * <0.05, ** <0.01 and *** <0.001 or exact FDR-value (if *FDR* < 0.10) vs. baseline. The exact *FDR*-values and fold changes of the proteins in each comparison are in Supporting information, Supplement 1.

For discovery science, *FDR* < 0.10 and/or unadjusted *p*-value approaches are typical in in human exercise proteomics studies (Deshmukh *et al*., 2021; Larsen *et al*., 2023; Roberts *et al*., 2024). Therefore, as we focus only on the 150 proteins altered by RT1 that may be retained, we next used a slightly more inclusive approach for the post-DT vs. baseline comparison (*FDR* < 0.10 in the whole proteome) while maintaining that the fold change still had to be greater than 1.2 (i.e. more than 20% increase). We identified 29 retained proteins (that included the four retained proteins identified above) to be still elevated after detraining following earlier training as well as after RT2 (vs. baseline, *FDR* < 0.10 and FC > 1.2) (Fig. 6C).

To gain further insight into biological processes, we conducted bioinformatics analysis of these retained proteins. The retained proteins were overrepresented especially in calcium-binding (Fig. 7A) and thus very differently than the reversible proteins involved in aerobic metabolism (Fig. 5D). GO term Molecular function showed that 7 out of 29 of the retained proteins have a role in calcium-binding (ANXA1, ANXA2, CAPN2, PLS3, S100A11, S100A13, S100A16). This is very different from 36 reversible proteins, of which only one had a molecular function in calcium-binding (SLC25A12). When analysing the 81 retained proteins using the SOM-approach, we found them being enriched in addition to calcium-binding also in actin filament binding. Indeed, 17 proteins of in total 81 proteins have GO:BP-term “actin”, “muscle contraction” or “myofibril” including several myosins/tropomyosin (ANXA1, ARPC2, ARPC3, CAMK2D, CASQ2, CORO1C, HSP90B1, IQGAP2, MARCKS, MYADM, MYH10, MYH9, MYL6B, PGM5, PLS3, TAGLN2, TPM4) and 6 proteins with term GO:BP term “cytoskeleton” (CLIC4, DPYSL2, IQGAP2, SEPTIN7, TUBA1C, TUBB2A).

**Figure 7.**
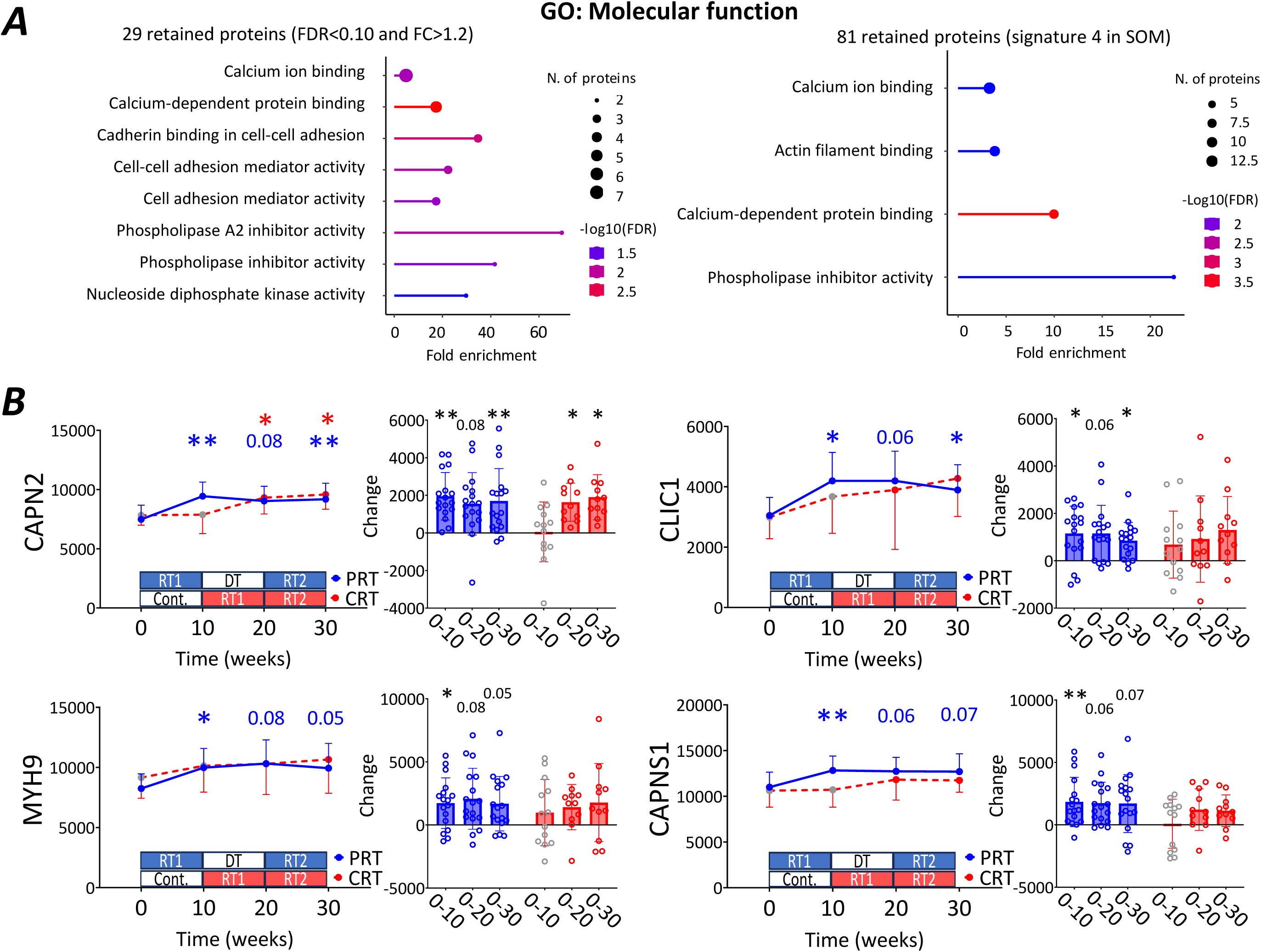
*A,* The 29 and 81 retained proteins (SOM) in GO have a molecular function especially in calcium binding. *B*, Previously described epigenetic memory genes CAPN2, CLIC1, and MYH9, as well as a regulatory subunit of CAPN2, CAPNS1, also demonstrate retained protein levels after DT. N=17 in PRT and n=11 in CRT, except n=13 in the first 10 weeks (control period). The proteins are plotted by applying the inverse log_2_ transformation to estimate their original mean ± SEM values. The statistics were conducted from log_2_ values. * <0.05, ** <0.01 or exact FDR-value (*FDR* < 0.10) vs. baseline. Figure *A* was extracted and modified from ShinyGO. The exact *FDR*-values and fold changes of the proteins in each comparison are in Supporting information, Supplement 1.

Next, we compared these 29 retained ‘memory’ proteins to previously published epigenetic memory genes, i.e. retained hypomethylation of the genes DNA after both RT and DT (Seaborne *et al*., 2018) and after HIIT training and detraining (Pilotto *et al*., 2024). We found that previously described epigenetic memory genes CAPN2, CLIC1, and MYH9, which are retained as hypomethylated, also demonstrate retained protein levels after DT following the earlier training in the present study. Of those proteins, CAPN2 and MYH9 were also identified to be hypomethylated genes following RT that also increased in 181 integrated skeletal muscle transcriptomes after RT (Turner et al., 2019). Of those two, CAPN2 had a similar time trend also in the CRT group, i.e. consistently increased as a result of RT with no change during the control period (Fig. 7B). CAPN2 is a catalytic subunit of calpain-2 forming a heterodimer complex with the small regulatory subunit (CAPNS1) to be a functional protease (Hyatt & Powers, 2020). In the present study CAPNS1 also demonstrates a retained protein profile after detraining (*FDR* = 0.0551) (Fig. 7B). Together, these findings suggest that calpain-2 is RT-induced "memory" protein, similar to CBR1 described above.

### The majority of the proteins decreased after RT1 have a reversible signature with one protein that is retained as down

Ten proteins decreased significantly after RT1 in the PRT group (Fig. 8A), of which almost all returned towards the baseline after 10 weeks of DT and were then decreased again after RT2 when compared to baseline (8 of 10 at *FDR* < 0.10 and 5 of 10 at *FDR* < 0.05). Interestingly, however, we identified one protein, Low density lipoprotein receptor-related protein-associated protein 1 (LRPAP1), to be slightly but consistently decreased at RT1 (*FDR* = 0.00531, DT (*FDR* = 0.0617) and RT2 (*FDR* < 0.001, Fig. 8B). Finally, after RT2 when compared to pre-RT2, two proteins decreased, MYH1 (IIX isoform, *FDR* = 0.0149) and ASAH1 (acid ceramidase, *FDR* = 0.0149) of which there was also similar magnitude (∼51 %, *FDR* = 0.0662) decrease in MYH1 following RT1 (Fig. 3E), as well in the CRT group (∼52 %, *FDR* = 0.138).

**Figure 8.**
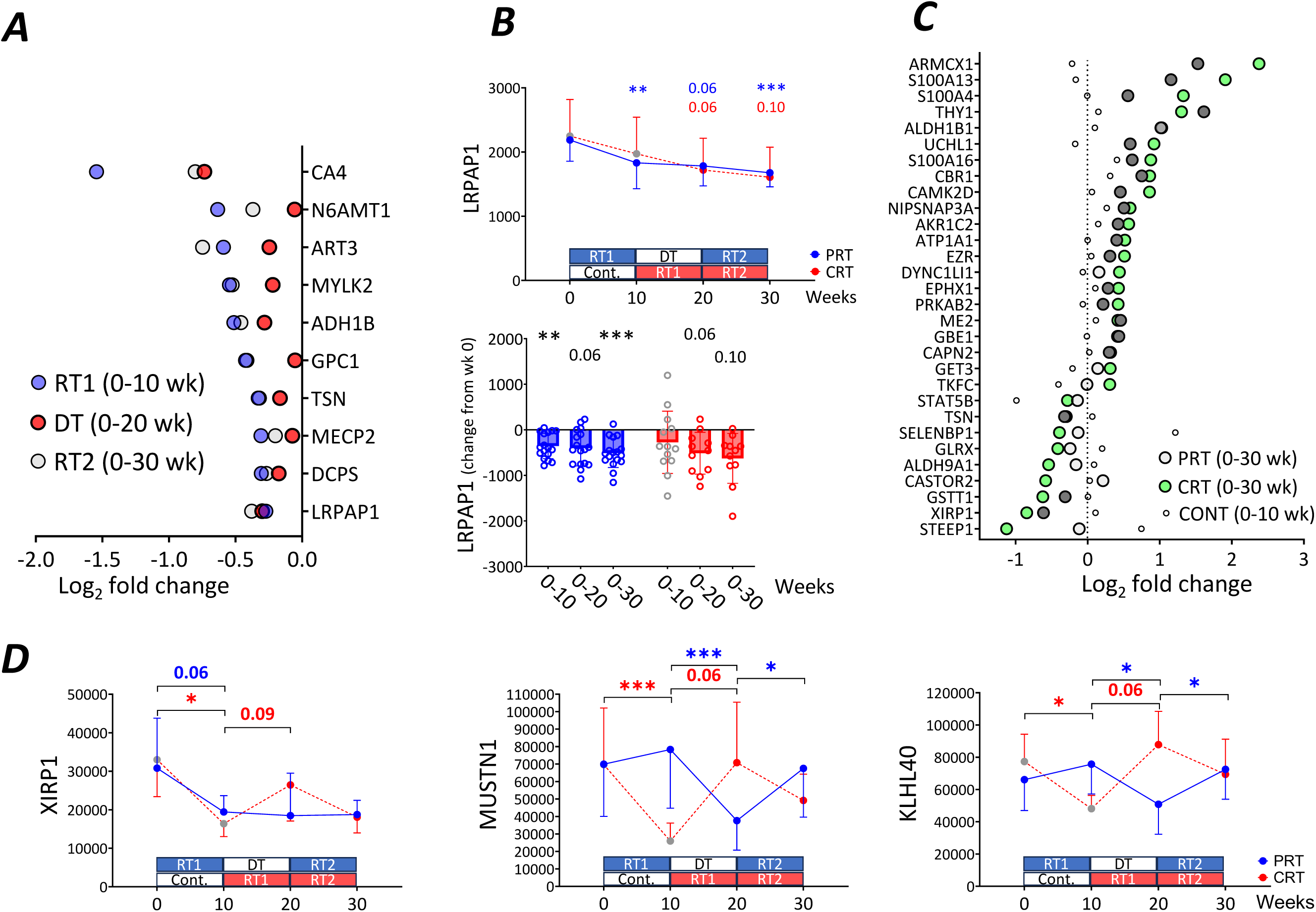
*A*, Ten proteins significantly decreased in response to RT1 in CRT group. *B*, LRPAP was a protein with decreased retained signature. *C*, Significantly altered proteins in CRT group (n=11) at 0 to 30 weeks plotted with the same proteins in the PRT (n=17) group and during the control period in the CRT group (n=13). Dark grey indicates *FDR* < 0.05 alteration also in the PRT group. *D*, Three proteins decreased during the control period possibly due to seasonal effect. The proteins are plotted by applying the inverse log_2_ transformation to estimate their original mean ± SEM values. The statistics were conducted from log_2_ values. * <0.05, ** <0.01 and *** <0.001 or exact FDR-value (if *FDR* < 0.10) vs. baseline. N=17 in PRT and n=11 in CRT, except n=13 in the first 10 weeks (control period). The exact *FDR*-values and fold changes of the proteins in each comparison are in Supporting information, Supplement 1.

### Outcomes of PRT when compared to CRT

As shown in Table 1 and earlier (Halonen *et al*., 2024), the eventual outcome (i.e. change from week 0 to 30) in muscle size or strength did not differ between PRT and CRT. Similarly, when comparing concurrent proteome adaptations, there were no significant between-group differences in the change (*FDR* > 0.2, Supporting information, Supplement 1). This suggests that the training break does not strongly affect skeletal muscle proteomic adaptation to 20 weeks of RT. Finally, to demonstrate similar responses in these two groups, the significantly altered proteins in the CRT group from 0-30 weeks behaved very similarly also in the PRT group during the same time-period (Fig. 8C).

### Potential seasonal changes in some skeletal muscle proteins

The three proteins altered during the no-training control period in the CRT group (Fig. 2A) included Xin Actin Binding Repeat Containing 1 (XIRP1), Musculoskeletal, Embryonic Nuclear Protein 1 (MUSTN1) and Kelch Like Family Member 40 (KLHL40). All these proteins demonstrated a decrease during the control period, perhaps due to a seasonal effect (from late winter to early summer) (Fig. 8D). For example, XIRP1 also simultaneously decreased in the PRT group. On the other hand, MUSTN1 and KLHL40 increased after RT2 in the PRT group but not after RT1 when their potential decreasing seasonal effect occurred.

## Discussion

In this study, we report substantial and reproducible remodelling of the human skeletal muscle proteome following repeated RT. Many of the proteins increasing after the first RT returned towards baseline after the training break while being increased again after a second period of RT. These proteins are especially involved in aerobic energy metabolism. However, several proteins induced by RT remained elevated even after ten weeks of detraining. The proteins with the retained profile include those involved in calcium binding, muscle contraction, and cytoskeleton. Herein, we are the first to display that human skeletal muscle possesses retained protein changes and a proteomic memory of RT-induced muscle growth.

Earlier epigenetic/transcriptomic studies in humans (Seaborne *et al*., 2018; Pilotto *et al*., 2024) and mice (Wen *et al*., 2021) have reported retention of epigenetic modifications after training (RT, HIIT, and weighted wheel running, respectively), which can lead to continued elevation of gene expression during detraining. The present study provides novel evidence to extend these results to the protein level. More specifically, of the individual proteins retained as altered after the 10-week training break, we identify especially calpain-2 (CAPN2), carbonyl reductase 1 (CBR1), and low-density lipoprotein receptor-related protein-associated protein (LRPAP1) as potential ‘muscle memory’ proteins. GO term Molecular function showed that, unlike reversible proteins, many of the retained proteins have a role in calcium-binding. One of the retained calcium-binding proteins was calpain-2 (CAPN2), which is also one of the few consistent hypomethylated muscle memory genes to RT (Seaborne *et al*., 2018; Turner *et al*., 2019) and HIIT (Pilotto *et al*., 2024). This gene transcript was also increased in 181 integrated skeletal muscle transcriptomes after RT (Turner *et al*., 2019) and was recently one of the most upregulated genes with a delay after a single bout of resistance exercise (Edman *et al*., 2024). The reliability of the result is also increased by the fact that this protein was consistently increased by RT in the present study and that its small regulatory subunit (CAPNS1) (Hyatt & Powers, 2020), also followed a similar retained protein signature. Recently, calpain-2 was also increased after 10 weeks of RT, especially post-exercise, and decreased already in 24 hours following a single exercise bout (Scarpelli et al., 2024), suggesting that it does not have a very long half-life in human skeletal muscle. Calpain-2 has an important role in the remodelling of skeletal muscle, such as essential myofibril disassembly during growth and atrophy stimuli, like exercise and inactivity, respectively (Hyatt & Powers, 2020). Therefore, more studies are needed to investigate both the effects of training and its cessation on calpain-2, and the physiological significance of these findings in skeletal muscle. When analysing the retained proteins using SOM-approach, we identified that many proteins were involved in muscle contraction or cytoskeleton in addition to calcium-binding. Thus, it appears that several proteins essential for muscle function have persisting levels in skeletal muscle following RT, which substantiates the effect on these pathways identified to be enriched in hypomethylation and gene expression after RT (Turner *et al*., 2019).

The overall RT-induced proteome involved processes such as chaperones and folding catalysis, protein processing in endoplasmic reticulum. These are well in line with a recent muscle proteomics study (Jessen *et al*., 2024) and studies showing that RT can induce endoplasmic reticulum (ER) stress and activate the unfolded protein response (UPR) pathways (Hentila *et al*., 2018). Intriguingly, we found that LRPAP1 was the only protein to decrease after RT that was also retained as decreased following detraining and the second RT period. LRPAP1 primarily functions as a molecular chaperone, but we are not aware of it being investigated in the context of exercise, skeletal muscle biology.

One RT-induced retained-up protein was enzyme carbonyl reductase (CBR1), which was also one of the most consistently RT-induced proteins. Carbonyl reductases belong to a class of oxidoreductase proteins catalysing the NADPH-dependent reduction of a large number of substrates (Forrest & Gonzalez, 2000). Transcript levels of CBR1 are significantly increased by RT based on MetaMEx (https://www.metamex.eu/) without an effect of inactivity (Pillon *et al*., 2020) and its protein levels were slightly (*FDR* < 0.10) elevated earlier after a shorter RT-intervention (Jessen *et al*., 2024). It is upregulated during skeletal muscle differentiation and regeneration through Nrf2 and appears to be an essential regulator of these processes in vitro through decreasing ROS levels and lipid peroxidation (Lim *et al*., 2013). This suggests that in addition to protecting muscle against damage caused by oxidative stress, it may be involved in the continuous remodelling of skeletal muscle following RT. More studies are needed to investigate the physiological significance of CBR1 in skeletal muscle.

Many proteins followed retained or semi-retained signatures following 10 weeks of training break. For many proteins, we expect that this reflects their long-term expression, i.e. potentially identifying them as muscle memory proteins. However, due to long half-lives in skeletal muscle (10 days on average in mice) (Rolfs *et al*., 2021) some of the protein elevations are probably retained during detraining, as speculated earlier after inactivity due to spaceflight and bed rest (Murgia *et al*., 2022). Nevertheless, we found several proteins decreasing back to baseline after detraining. These reversible proteins were particularly enriched in energy metabolism, mitochondrial function, redox homeostasis, and especially the TCA cycle and electron transport chain. This aligns with the known short *in-vivo* half-life of enzymes and fast decrease of aerobic capacity after detraining (Mujika & Padilla, 2000) and downregulated mitochondrial proteome after muscle deloading in low gravity (Murgia *et al*., 2022). One example of a reversible RT-upregulated protein essential in skeletal muscle energy metabolism was enzyme NAMPT. It has been previously shown to be upregulated by RT (de Guia *et al*., 2019; Lamb *et al*., 2020), and we expand these findings by showing that the effect is reversible and repeatable upon later retraining.

This is the first study to investigate in a longitudinal (within-individual) design whether the proteome responses to exercise are systematic. Our study also included a control period to exclude results due to, for example, seasonal changes or other random effects. Our data shows a much more robust effect of RT than the control period on proteomics, but we speculate that also potential seasonal changes may affect some muscle proteins as seasonal changes in gene expression are known to exist in human cells (Dopico *et al*., 2015). The three proteins altered during the no-training control period in the CRT group included MUSTN1, XIRP1, and KLHL40. These proteins demonstrated a decrease during the control period, perhaps due to a seasonal effect (from late winter to early summer). For example, XIRP1 also simultaneously decreased in the PRT group. On the other hand, MUSTN1 and KLHL40 increased after RT2 in the PRT group but not after RT1 when their potential decreasing seasonal effect occurred. MUSTN1 is a protein recently shown to be a secreted protein in skeletal muscle, decreased after hindlimb unloading and rescued after reloading (Ducommun *et al*., 2024). We expand these findings by showing that RT also increases this protein in humans. Another kelch-like family protein KLHL41 recently identified as an RT- induced protein (Jessen *et al*., 2024) did not change during the control period, but was consistently responsive to both RT1 and RT2. Interestingly, in addition to very similar time courses in both PRT and CRT groups, RT-induced changes in MUSTN1 correlated strongly and consistently with changes in KLHL40 (RT1 vs. baseline, RT2 vs. baseline and RT2 vs. DT: r = 0.70 - 0.80 and *P* < 0.002). Moreover, when the changes in *MUSTN1* were correlated with all other gene expression changes, *KLHL40* emerged as the strongest link also in MetaMEx (https://www.metamex.eu/) (Pillon et al., 2020). This suggests MUSTN1 and KLHL40 may have common upstream regulation, which should be investigated mechanistically in the future.

We report in our periodic RT group reproducible protein changes after the first RT and the whole period, as two thirds of the RT1-induced proteins were also induced after the second RT period when compared to the baseline. Moreover, when the second RT period was compared to its pre-state (i.e. to post-detraining), more than half of the proteins altered behaved very similarly after RT1 and, overall, significantly correlated with the responses in RT1. Based on fold change (with *FDR* < 0.05), the top upregulated proteins after the first RT period were Quinone oxidoreductase PIG3 (TP53I3), Thrombospondin-4 (THBS4), Adenylate kinase 4 (AK4), Thy-1 membrane glycoprotein (THY1) and Ras GTPase-activating-like protein (IQGAP2), and the most decreased protein was Carbonic anhydrase 4 (CA4). Based on FDR-value, the most consistently altered (increased) protein after the first RT period was the previously mentioned retained protein CBR1 (*FDR* = 0.0000586). When compared to baseline, more proteins were altered after the second RT period than the first, and many of those proteins were enriched specifically in aerobic metabolism pathways and biological processes, which did not appear to occur in the continuous training group. This suggests that some proteins have greater levels after the second period of RT, but only when there is a break between the training periods.

Finally, we also compared our data to a recently published four-week RT intervention (Jessen *et al*., 2024). Of the 140 proteins increased in the present study, about half (71) were identified. Of those proteins, 24 were also increased in (Jessen *et al*., 2024), meaning a ∼34 % overlap. We also found many similar protein changes, such as increases in KLHL40 and KLHL41, of which the latter was mechanistically shown to regulate muscle cell size *in vitro* (Jessen *et al*., 2024). Overall, these studies suggest a considerable reproducibility of the effect of RT on muscle proteome.

The future human RT proteomics studies focusing on muscle memory at the protein level should investigate the time-course of proteins following RT and DT. Excluding the TDR group in the middle of the 20-week RT period, we obtained the post-RT muscle biopsies ∼6 days after the last exercise bout, which is expected to be sufficient to minimize the acute effects of the last resistance exercise bout as the participants were already well accustomed to the RT (Egan & Sharples, 2023). Nevertheless, the future studies should also investigate what happens to the muscle proteome acutely after single bout of resistance exercise and whether this response is altered after detraining and retraining. Moreover, different types of research designs are needed to potentially allow to distinguish proteomic memory of muscle growth from the proteomic memory or signature of RT *per se*. In addition, combining proteomics approaches to metabolomics, transcriptomics, and epigenetics as well as other epigenetic modifications such as chromatin accessibly via ATAC-seq in the future, are crucial to advance our understanding of muscle memory. Finally, the physiological significance of long-term protein changes and proteomic memory of RT-induced muscle growth should be investigated in the future by mechanistic experiments.

In summary, we have presented the first in-depth data on proteomic remodelling to resistance training, followed by a training break and retraining. We first demonstrate that the proteome adaptations to resistance training are reproducible. We further identify proteins that either return to baseline or remain elevated even after 2.5 months without training. This data supports the presumption that human skeletal muscle possesses a potential proteomic memory of resistance training-induced muscle growth following resistance training.

## ACKNOWLEDGEMENTS

The authors would like to thank Antti Tuhkala, Erik Niemi, MD Milla Kelahaara, and the other laboratory personnel, students, research assistants, and participants for their time and effort in completing this study.

## DATA AVAILABILITY STATEMENT

The data that supports the findings of this study are available from the corresponding author upon reasonable request.

## References

Ahtiainen JP, Hoffren M, Hulmi JJ, Pietikäinen M, Mero AA, Avela J & Häkkinen K (2010). Panoramic ultrasonography is a valid method to measure changes in skeletal muscle cross-sectional area. Eur J Appl Physiol 108, 273–279.

Beltrà M, Pöllänen N, Fornelli C, Tonttila K, Hsu MY, Zampieri S, Moletta L, Corrà S, Porporato PE, Kivelä R, Viscomi C, Sandri M, Hulmi JJ, Sartori R, Pirinen E & Penna F (2023). NAD+ repletion with niacin counteracts cancer cachexia. Nat Commun 14, 1849.

Blocquiaux S, Ramaekers M, Van Thienen R, Nielens H, Delecluse C, De Bock K & Thomis M (2022). Recurrent training rejuvenates and enhances transcriptome and methylome responses in young and older human muscle. JCSM Rapid Commun 5, 10–32.

Cumming KT, Reitzner SM, Hanslien M, Skilnand K, Seynnes OR, Horwath O, Psilander N, Sundberg CJ & Raastad T (2024). Muscle memory in humans: evidence for myonuclear permanence and longLterm transcriptional regulation after strength training. J Physiol 602, 4171–4193.

Demichev V, Messner CB, Vernardis SI, Lilley KS & Ralser M (2020). DIA-NN: neural networks and interference correction enable deep proteome coverage in high throughput. Nat Methods 17, 41–44.

Demichev V, Szyrwiel L, Yu F, Teo GC, Rosenberger G, Niewienda A, Ludwig D, Decker J, Kaspar-Schoenefeld S, Lilley KS, Mülleder M, Nesvizhskii AI & Ralser M (2022). dia-PASEF data analysis using FragPipe and DIA-NN for deep proteomics of low sample amounts. Nat Commun 13, 3944.

Deshmukh AS, Steenberg DE, Hostrup M, Birk JB, Larsen JK, Santos A, Kjøbsted R, Hingst JR, Schéele CC, Murgia M, Kiens B, Richter EA, Mann M & Wojtaszewski JFP (2021). Deep muscle-proteomic analysis of freeze-dried human muscle biopsies reveals fiber type-specific adaptations to exercise training. Nat Commun 12, 304.

Dopico XC, Evangelou M, Ferreira RC, Guo H, Pekalski ML, Smyth DJ, Cooper N, Burren OS, Fulford AJ, Hennig BJ, Prentice AM, Ziegler A-G, Bonifacio E, Wallace C & Todd JA (2015). Widespread seasonal gene expression reveals annual differences in human immunity and physiology. Nat Commun 6, 7000.

Du J, Yun H, Wang H, Bai X, Su Y, Ge X, Wang Y, Gu B, Zhao L, Yu J-G & Song Y (2024). Proteomic Profiling of Muscular Adaptations to Short-Term Concentric Versus Eccentric Exercise Training in Humans. Molecular & Cellular Proteomics 23, 100748.

Ducommun S et al. (2024). Mustn1 is a smooth muscle cell-secreted microprotein that modulates skeletal muscle extracellular matrix composition. Mol Metab 82, 101912.

Edman S et al. (2024). The 24-hour molecular landscape after exercise in humans reveals MYC is sufficient for muscle growth. EMBO Rep; DOI: 10.1038/s44319-024-00299-z.

Eftestøl E, Ochi E, Juvkam IS, Hansson K & Gundersen K (2022). A juvenile climbing exercise establishes a muscle memory boosting the effects of exercise in adult rats. Acta Physiologica; DOI: 10.1111/apha.13879.

Egan B & Sharples AP (2023). Molecular responses to acute exercise and their relevance for adaptations in skeletal muscle to exercise training. Physiol Rev 103, 2057–2170.

Egner IM, Bruusgaard JC, Eftestøl E & Gundersen K (2013). A cellular memory mechanism aids overload hypertrophy in muscle long after an episodic exposure to anabolic steroids. J Physiol 591, 6221–6230.

Forrest GL & Gonzalez B (2000). Carbonyl reductase. Chem Biol Interact 129, 21–40.

Ge SX, Jung D & Yao R (2020). ShinyGO: a graphical gene-set enrichment tool for animals and plants. Bioinformatics 36, 2628–2629.

de Guia RM, Agerholm M, Nielsen TS, Consitt LA, Søgaard D, Helge JW, Larsen S, Brandauer J, Houmard JA & Treebak JT (2019). Aerobic and resistance exercise training reverses ageLdependent decline in NAD ^+^ salvage capacity in human skeletal muscle. Physiol Rep; DOI: 10.14814/phy2.14139.

Halonen EJ, Gabriel I, Kelahaara MM, Ahtiainen JP & Hulmi JJ (2024). Does Taking a Break Matter—Adaptations in Muscle Strength and Size Between Continuous and Periodic Resistance Training. Scand J Med Sci Sports; DOI: 10.1111/sms.14739.

Hentila J, Ahtiainen JP, Paulsen G, Raastad T, Hakkinen K, Mero AA & Hulmi JJ (2018). Autophagy is induced by resistance exercise in young men, but unfolded protein response is induced regardless of age. Acta Physiol (Oxf) 224, e13069.

Hulmi JJ, Tannerstedt J, Selänne H, Kainulainen H, Kovanen V & Mero AA (2009). Resistance exercise with whey protein ingestion affects mTOR signaling pathway and myostatin in men. J Appl Physiol; DOI: 10.1152/japplphysiol.00087.2009.

Hyatt HW & Powers SK (2020). The Role of Calpains in Skeletal Muscle Remodeling with Exercise and Inactivity-induced Atrophy. Int J Sports Med 41, 994–1008.

Jassal B et al. (2019). The reactome pathway knowledgebase. Nucleic Acids Res; DOI: 10.1093/nar/gkz1031.

Jessen S, Quesada JP, Di Credico A, MorenoLJusticia R, Wilson R, Jacobson G, Bangsbo J, Deshmukh AS & Hostrup M (2024). Beta _2_ LAdrenergic Stimulation Induces Resistance TrainingLLike Adaptations in Human Skeletal Muscle: Potential Role of KLHL41. Scand J Med Sci Sports; DOI: 10.1111/sms.14736.

Lamb DA, Moore JH, Mesquita PHC, Smith MA, Vann CG, Osburn SC, Fox CD, Lopez HL, Ziegenfuss TN, Huggins KW, Goodlett MD, Fruge AD, Kavazis AN, Young KC & Roberts MD (2020). Resistance training increases muscle NAD+ and NADH concentrations as well as NAMPT protein levels and global sirtuin activity in middle-aged, overweight, untrained individuals. Aging 12, 9447–9460.

Larsen JK, Kruse R, Sahebekhtiari N, Moreno-Justicia R, Gomez Jorba G, Petersen MH, de Almeida ME, Ørtenblad N, Deshmukh AS & Højlund K (2023). High-throughput proteomics uncovers exercise training and type 2 diabetes–induced changes in human white adipose tissue. Sci Adv; DOI: 10.1126/sciadv.adi7548.

Liebermeister W, Noor E, Flamholz A, Davidi D, Bernhardt J & Milo R (2014). Visual account of protein investment in cellular functions. Proceedings of the National Academy of Sciences 111, 8488–8493.

Lim S, Shin JY, Jo A, K.R J, Nguyen MN, Choi TG, Kim J, Park J-H, Eun YG, Yoon K-S, Ha J & Kim SS (2013). Carbonyl reductase 1 is an essential regulator of skeletal muscle differentiation and regeneration. Int J Biochem Cell Biol 45, 1784–1793.

Meier F, Brunner A-D, Frank M, Ha A, Bludau I, Voytik E, Kaspar-Schoenefeld S, Lubeck M, Raether O, Bache N, Aebersold R, Collins BC, Röst HL & Mann M (2020). diaPASEF: parallel accumulation-serial fragmentation combined with data-independent acquisition. Nat Methods 17, 1229–1236.

Meier F, Brunner A-D, Koch S, Koch H, Lubeck M, Krause M, Goedecke N, Decker J, Kosinski T, Park MA, Bache N, Hoerning O, Cox J, Räther O & Mann M (2018). Online Parallel Accumulation-Serial Fragmentation (PASEF) with a Novel Trapped Ion Mobility Mass Spectrometer. Mol Cell Proteomics 17, 2534–2545.

Moesgaard L, Moreno-Justicia R, Schmalbruch J, Jessen S, Stocks B, Bangsbo J, Deshmukh AS & Hostrup M (2024). Muscle fiber proteomics reveals sex-and fiber type-specific adaptations to resistance training. ; DOI: 10.1101/2024.09.16.612737.

Mujika I & Padilla S (2000). Detraining: Loss of Training-Induced Physiological and Performance Adaptations. Part I. Sports Medicine 30, 79–87.

Murach KA, Mobley CB, Zdunek CJ, Frick KK, Jones SR, McCarthy JJ, Peterson CA & Dungan CM (2020). Muscle memory: myonuclear accretion, maintenance, morphology, and miRNA levels with training and detraining in adult mice. J Cachexia Sarcopenia Muscle 11, 1705– 1722.

Murgia M, Ciciliot S, Nagaraj N, Reggiani C, Schiaffino S, Franchi M V, Pišot R, Šimunič B, Toniolo L, Blaauw B, Sandri M, Biolo G, Flück M, Narici M V & Mann M (2022). Signatures of muscle disuse in spaceflight and bed rest revealed by single muscle fiber proteomics. PNAS Nexus; DOI: 10.1093/pnasnexus/pgac086.

Pillon NJ, Gabriel BM, Dollet L, Smith JAB, Sardon Puig L, Botella J, Bishop DJ, Krook A & Zierath JR (2020). Transcriptomic profiling of skeletal muscle adaptations to exercise and inactivity. Nat Commun 11, 470-w.

Pilotto AM, Turner DC, Mazzolari R, Crea E, Brocca L, Pellegrino MA, Miotti D, Bottinelli R, Sharples AP & Porcelli S (2024). Human skeletal muscle possesses an epigenetic memory of high intensity interval training. American Journal of Physiology-Cell Physiology; DOI: 10.1152/ajpcell.00423.2024.

Psilander N, Eftestøl E, Cumming KT, Juvkam I, Ekblom MM, Sunding K, Wernbom M, Holmberg H-C, Ekblom B, Bruusgaard JC, Raastad T & Gundersen K (2019). Effects of training, detraining, and retraining on strength, hypertrophy, and myonuclear number in human skeletal muscle. J Appl Physiol 126, 1636–1645.

Roberts MD, Ruple BA, Godwin JS, McIntosh MC, Chen S-Y, Kontos NJ, Agyin-Birikorang A, Michel M, Plotkin DL, Mattingly ML, Mobley B, Ziegenfuss TN, Fruge AD & Kavazis AN (2024). A novel deep proteomic approach in human skeletal muscle unveils distinct molecular signatures affected by aging and resistance training. Aging; DOI: 10.18632/aging.205751.

Robinson MM, Dasari S, Konopka AR, Johnson ML, Manjunatha S, Esponda RR, Carter RE, Lanza IR & Nair KS (2017). Enhanced Protein Translation Underlies Improved Metabolic and Physical Adaptations to Different Exercise Training Modes in Young and Old Humans. Cell Metab 25, 581–592.

Rolfs Z, Frey BL, Shi X, Kawai Y, Smith LM & Welham N V. (2021). An atlas of protein turnover rates in mouse tissues. Nat Commun 12, 6778.

Scarpelli MC, Bergamasco JGA, Godwin JS, Mesquita PHC, Chaves TS, Silva DG, Bittencourt D, Dias NF, Medalha Junior RA, Carello Filho PC, Angleri V, Costa LAR, Kavazis AN, Ugrinowitsch C, Roberts MD & Libardi CA (2024). Resistance training-induced changes in muscle proteolysis and extracellular matrix remodeling biomarkers in the untrained and trained states. Eur J Appl Physiol 124, 2749–2762.

Seabold S & Perktold J (2010). Statsmodels: Econometric and Statistical Modeling with Python.

Seaborne RA, Strauss J, Cocks M, Shepherd S, O’Brien TD, van Someren KA, Bell PG, Murgatroyd C, Morton JP, Stewart CE & Sharples AP (2018). Human Skeletal Muscle Possesses an Epigenetic Memory of Hypertrophy. Sci Rep **8**, 1893–1898.

Sharples AP & Turner DC (2023). Skeletal muscle memory. American Journal of Physiology-Cell Physiology 324, C1274–C1294.

Snijders T, Leenders M, de Groot LCPGM, van Loon LJC & Verdijk LB (2019). Muscle mass and strength gains following 6 months of resistance type exercise training are only partly preserved within one year with autonomous exercise continuation in older adults. Exp Gerontol 121, 71–78.

Timmons JA, Szkop KJ & Gallagher IJ (2015). Multiple sources of bias confound functional enrichment analysis of global -omics data. Genome Biol 16, 186.

Turner DC et al. (2020). DNA methylation across the genome in aged human skeletal muscle tissue and muscle-derived cells: the role of HOX genes and physical activity. Sci Rep 10, 15360-z.

Turner DC, Seaborne RA & Sharples AP (2019). Comparative Transcriptome and Methylome Analysis in Human Skeletal Muscle Anabolism, Hypertrophy and Epigenetic Memory. Sci Rep 9, 4251.

Virtanen P et al. (2020). SciPy 1.0: fundamental algorithms for scientific computing in Python. Nat Methods 17, 261–272.

Wen Y, Dungan CM, Mobley CB, Valentino T, von Walden F & Murach KA (2021). Nucleus Type-Specific DNA Methylomics Reveals Epigenetic “Memory” of Prior Adaptation in Skeletal Muscle. Function; DOI: 10.1093/function/zqab038.

Wijesooriya K, Jadaan SA, Perera KL, Kaur T & Ziemann M (2022). Urgent need for consistent standards in functional enrichment analysis. PLoS Comput Biol 18, e1009935.

